# Host transcriptional signatures associated with disease tolerance and environmental persistence in a mosquito–microsporidian system

**DOI:** 10.1101/2024.09.22.613703

**Authors:** Luís M. Silva

## Abstract

Despite strong theory linking parasite life history to virulence and transmission, how parasite strategies affect host responses remains poorly understood. Here, experimental evolution and RNA sequencing were combined in the mosquito–microsporidian system *Anopheles gambiae–Vavraia* culicis to test whether parasite lineages selected *for* early or *late* transmission differ in the host responses they elicit. *Early* lineages were selected from hosts that died early in infection, whereas *late* lineages were selected from hosts that died later, generating parasite populations that differ in growth, virulence and environmental persistence. Mosquitoes were infected with evolved parasite lineages and sampled at a common sporulating stage. A *reference* infection response was defined relative to *uninfected* controls. *Early*- and *late*-selected parasites showed consistent, yet partly distinct, changes in host gene expression. Co-expression network analysis identified gene modules whose activity covaried with parasite traits, including virulence and environmental persistence. One module enriched for tissue maintenance and metabolic functions was negatively associated with virulence, consistent with reduced damage. Another module enriched for membrane and signalling functions was positively associated with environmental persistence, suggesting that host state at transmission may influence the environment in which transmission stages develop. Together, these results identify distinct host states linked to parasite life-history variation.

## Introduction

Parasites have a diverse array of strategies and life histories, shaped by selection acting across their infection cycle: from growth within hosts to transmission between them [1–4]. Virulence, defined as the reduction in host fitness caused by infection, arises from interactions between parasite traits and host responses rather than being a fixed property of the parasite alone [5,6]. Hosts can reduce the costs of infection by limiting parasite load (resistance) or by limiting the damage caused by infection or the immune response to it (tolerance) [7–10]. Tolerance can involve damage repair, metabolic reallocation, or maintenance of barrier integrity, and is therefore expected to elicit different host responses than resistance, which is centred on parasite killing [11–17]. Parasites, in turn, can increase virulence either by increasing their within-host density (greater host exploitation) or by increasing the damage caused per parasite unit through the production of pathogenic factors (per-parasite pathogenicity; for instance, toxins) [1,6,7,18]. Classical theory often compresses this complexity into a single trade-off between transmission rate and virulence [19,20], yet empirical work shows that such simplified expectations capture only part of how virulence evolves, especially when different stages of the infection cycle are under selection [3,21–23].

Many parasites must allocate resources between growth within hosts and the production of asexual and sexual transmission stages, which can be released into the environment or transmitted via vectors (e.g., *Plasmodium spp.*) [2,4,23,24]. This may create trade-offs between within-host growth and the production and persistence of transmission stages, which can influence virulence [2,25,26]. At the same time, infections occur within hosts with their own life histories [27–32]. When infection shortens expected lifespan, hosts may shift investment from somatic maintenance to early reproduction, a response known as terminal investment [33–35]. These interactions among parasite life history, host life history, and modes of transmission shape infection outcomes, yet the host responses underlying resistance, tolerance, and transmission remain poorly understood, particularly in vector hosts. Gene expression data enable the identification of host states associated with these processes and link parasite strategies to host biology. In vector systems, this overlap is especially relevant because parasite transmission depends on host survival long enough to complete the life cycle.

The microsporidian parasite *Vavraia culicis* [36] and the mosquito host *Anopheles gambiae* provide a natural model to study how parasite life-history variation is reflected in host responses and consequent transmission. *V. culicis* is an obligate intracellular parasite that infects mosquito larvae, proliferates through larval and adult stages, and is mainly transmitted when infected adults die and release spores into water [36,37]. Parasites must allocate resources between growth within mosquitoes and the production of transmission stages that remain infective outside the host. In vector systems, host tissues that parasites must cross or exploit can act as filters that shape transmission. For instance, in *Plasmodium*, the mosquito midgut epithelium and complement-like factor TEP1 eliminate a large fraction of invading parasites, making this barrier a key determinant of transmission success [38]. Similar constraints have been described for arboviruses at the level of midgut and salivary-gland barriers [33,34]. However, comparatively little is known about how parasite strategies associated with environmental transmission are reflected in host responses. Similar constraints have been described for arboviruses at the level of midgut and salivary-gland barriers [39,40]. Host genes and molecular processes that limit parasite development and transmission are increasingly well characterized in vector-borne systems [41–45]. In contrast, host determinants remain poorly understood for parasites with environmental transmission, despite theoretical predictions that host traits and environmental survival can jointly shape transmission and virulence [2,24,26,46].

Experimental evolution of *V. culicis in An. gambiae* has shown that parasite life history can evolve rapidly under selection on the timing of transmission [1,47,48]. Parasite lineages selected from hosts that died early in infection (hereafter, *early*) and from hosts that died later (*late*) were derived from an unselected *reference* population and selected for multiple generations [1]. Selection on transmission timing does not directly select for virulence, and *early* and *late* regimes do not correspond to the timing of mortality they induce. Instead, they impose different constraints on parasite growth and transmission. As a result, *late* lines showed higher exploitation, characterized by faster spore growth and enhanced use of host resources, including energy reserves and iron, leading to higher parasite burdens and virulence, whereas *early* lines showed lower exploitation but produced spores that remained infective for longer outside the host [1,47,48]. These parasite lineages also differed in the host life-history responses they induced, including development time, timing of reproduction, and allocation between reproduction and somatic maintenance [1]. Together, these findings show that selection on transmission timing reshapes parasite exploitation, virulence, environmental persistence, and host life history. However, they do not reveal how these differences are reflected in host gene expression.

Here, experimental evolution and RNA sequencing (RNA-seq) were combined to test whether parasite lineages that differ in virulence, exploitation, and environmental persistence are associated with distinct host transcriptional responses. Mosquitoes were infected with independently evolved parasite lineages and sampled at a common sporulating stage of infection (day 10 of adulthood), allowing host gene expression to be compared across parasite strategies while controlling for infection stage. In this host-parasite system, transmission occurs via the environment, providing a way to examine how host processes during infection are linked to the production and release of transmission stages. Because *early* and *late* lineages differ in growth, exploitation, and environmental persistence, distinct host transcriptional states were expected, particularly in processes related to resource use and tissue maintenance. A *reference* infection response was first defined relative to *uninfected* controls. *Early* and *late* infections were then compared to identify shared and regime-specific changes in gene expression. Co-expression network analysis (WGCNA) was used to identify gene modules whose activity covaried with infection background and parasite traits, including virulence and environmental persistence. These analyses show that parasite life-history variation is reflected in distinct host transcriptional states aligned with differences in virulence and environmental persistence, highlighting the role of host condition during infection in shaping transmission outcomes.

## Results

Mosquito transcriptomes were compared across infections with experimentally evolved microsporidian parasites to test how microsporidian infection strategies are associated with host responses and how these responses relate to virulence, parasite load, parasite environmental persistence and host fecundity. A single *An. gambiae* (Kisumu) background was either left *uninfected*, infected with a *reference V. culicis* population, or infected with parasite lines that had been experimentally evolved under contrasting selection on transmission timing [26,27] **(Fig. 1a)**. These selection regimes generated consistent differences among parasite lines: *late* lines reached higher parasite loads, were more virulent, and showed reduced environmental persistence, whereas *early* lines reached lower loads and virulence but produced more persistent transmission stages [1,48]. Five independently evolved parasite lineages were available within each of the *early* and *late* regimes, providing replication for evolutionary inference. In contrast, a single *reference* and a single *uninfected* lineage served as physiological anchors **(Fig. 1b, c)**.

**Figure 1.**
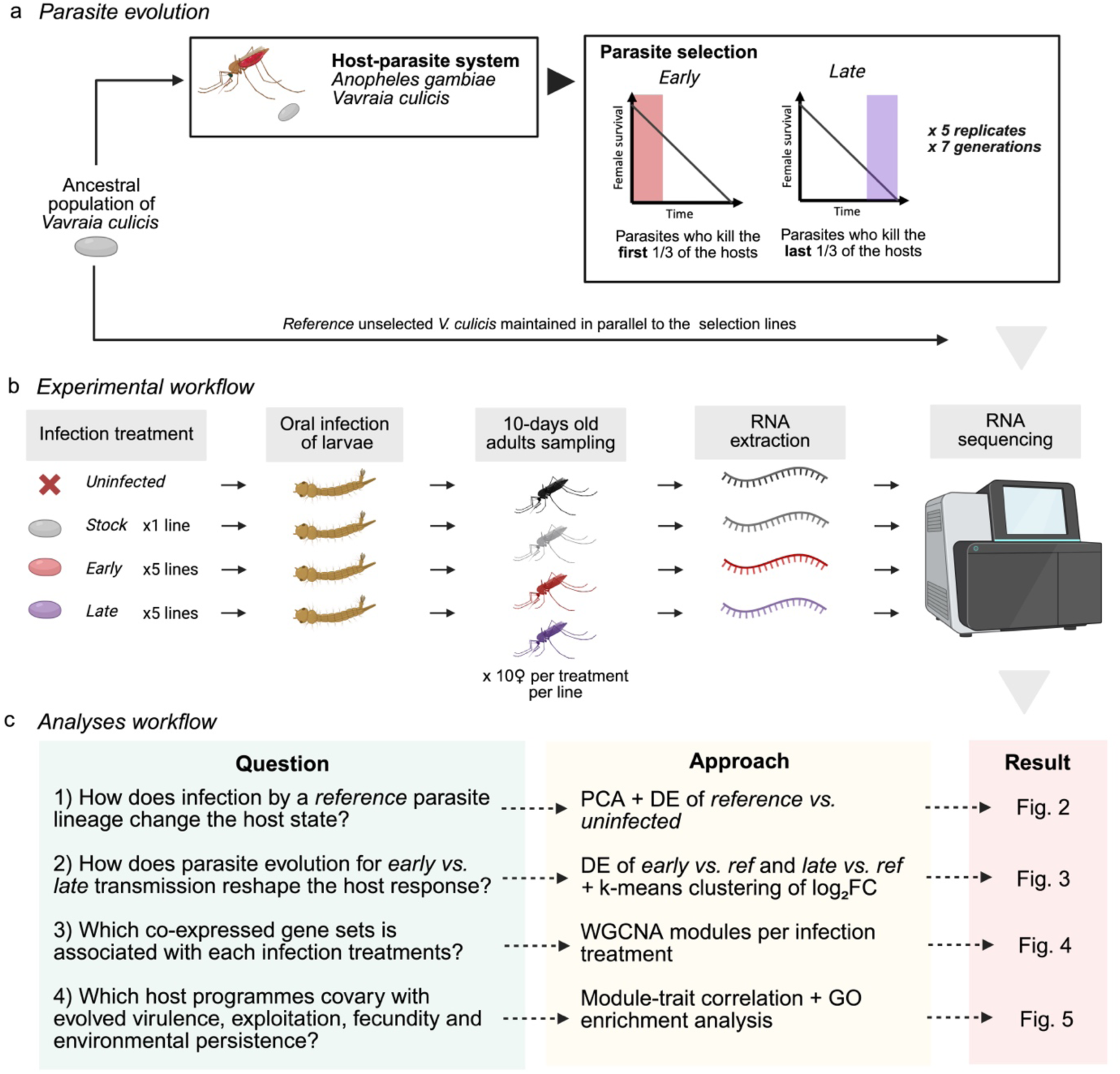
Parasite evolution, experimental and analytical workflow. **(a)** An ancestral *Vavraia culicis* population infecting *Anopheles gambiae* was split into selection regimes. In the *early* regime, spores were collected from hosts dying in the first third of the survival distribution; in the *late* regime, spores were collected from hosts dying in the last third. Selection was applied for seven generations in five replicate parasite lineages per regime, while an unselected *reference* population was maintained in parallel. **(b)** Larval *An. gambiae* were orally exposed to either no spores (*uninfected*), the *reference* parasite (1 lineage), or *early* or *late* parasites (5 lineages each). Adult females were sampled at day 10 post-emergence, when infections were chronic and sporulating, and RNA was extracted from four independent pools of 10 females per lineage for RNA sequencing. **(c)** Four main questions were addressed using different analytical approaches: the core host response to *reference* infection (PCA and differential expression, Fig. 2), divergence in host responses to *early vs. late* parasites (Differentially Expression [DE] and k-means clustering of log₂ fold-changes, Fig. 3), co-expressed gene sets associated with infection backgrounds (WGCNA modules, Fig. 4), and host programmes covarying with evolved virulence, host exploitation and fecundity, and parasite environmental persistence (WGCNA module–trait correlations, GO enrichment and hub-gene expression, Figs. 4–5).

Larval mosquitoes were exposed to the different parasite lineages using a standardized spore dose across treatments, and adult females were sampled at a common time point when infections were mature (10 days of adulthood). Previous work on these lines had demonstrated clear differences in virulence (measured as host mortality), host exploitation (parasite growth and resource use), within-host parasite growth (parasite load dynamics), fecundity, and environmental persistence [1,47,48]. RNA was extracted from pools of ten females per sample. Each lineage (and the *uninfected* control) was represented by four replicate pools, yielding 44 transcriptomes spanning infection treatments and selection regimes.

Across the twelve lineages (one *uninfected*, one *reference*, five *early*, five *late*), 20–38 million reads were obtained per sample, covering over 9,900 genes **(Supplementary Table 1 and Supplementary Fig. 1)**. Reads aligned successfully to the *An. gambiae* genome, and standard quality checks confirmed that libraries and mapping were suitable for downstream analyses **(Supplementary Fig. 2 and 3)**.

### Core response to infection

The *reference* infection induced a strong and coherent shift in mosquito gene expression relative to *uninfected* controls. In a PCA restricted to *reference* and *uninfected* samples, PC1 explained a significant proportion of the variance and separated samples almost entirely by infection status **(Fig. 2a; Supplementary Table 2)**. PC2 captured variation among replicate pools within treatments, indicating some heterogeneity among samples, but this variation did not obscure the clear separation between infected and uninfected groups.

**Figure 2.**
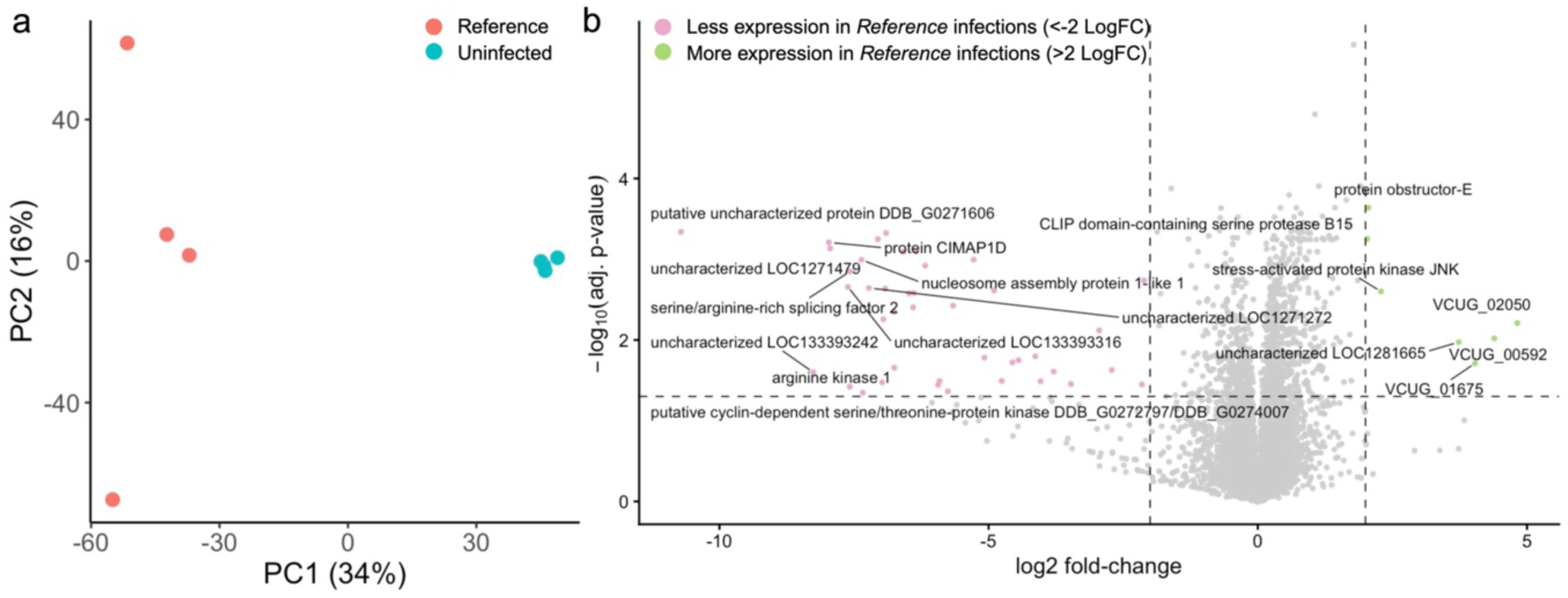
Host transcriptional response to a *Reference* infection. **(a)** Principal component analysis (PCA) of normalized gene expression in mosquitoes exposed to the *reference V. culicis* line (*reference*, **red**) or left *uninfected* (*uninfected*, **blue**). Each point represents one pool of ten females (four independent replicate pools per treatment). PCA was performed on the centred and scaled expression matrix after retaining the 75% most variable genes, a commonly used approach to focus on genes contributing most to variation in transcriptomic datasets. The axes show the percentage of variance explained by PC1 and PC2. *Reference* infections separated clearly from *uninfected* controls along PC1, indicating a strong and consistent infection signature. **(b)** Volcano plot of gene expression differences between *reference* and *uninfected* treatments. For each gene, the log₂ fold-change (*reference* - *uninfected*) and the –log₁₀ of the Benjamini–Hochberg adjusted *p*-value are shown. Genes with FDR < 0.05 and |log₂FC| > 2 are coloured according to whether they show lower (**pink**) or higher (**green**) expression in *Reference* infections. The genes with a larger effect size in each direction are labelled.

Differential expression (DE) analysis between *reference* and *uninfected* treatments confirmed that infection altered the expression of many genes **(Fig. 2b)**. Hundreds of genes showed significant differences (false discovery rate, FDR < 0.05; |log₂ fold change| > 2), with both strongly up-regulated and down-regulated loci. Up-regulated genes included several known or putative immune effectors and stress response genes, consistent with activation of canonical defence pathways. Down-regulated genes were enriched for more general metabolic and homeostatic functions, suggesting that infection also involves a reorganization of baseline physiology. This core infection signature was therefore used as a reference state against which evolved parasite infections were compared.

### Evolved parasite strategies modify the core response

When only infected samples (*reference*, *early,* and *late*) were considered, the PCA showed separation among infection regimes, with samples clustering by parasite lineage, and replicate pools from the same lineage grouping together **(Fig. 3a; Supplementary Table 3, and Supplementary Fig. 4)**. In particular, samples corresponding to the same independent parasite lineage (1–5) grouped together irrespective of the selection regime. Lineage 4 of the *late* treatment was separated from other samples along PC1, whereas PC2 separated lineage groups across treatments. This pattern indicates that a substantial proportion of the variance in the dataset is structured by lineage identity. Such variation is consistent with independently evolving lineages following partially distinct trajectories under the same selection regime. However, the contribution of lineage-level effects cannot be resolved from this analysis alone. Evolutionary inference in this design, therefore, relies on consistent differences between *early* and *late* treatments across independent lineages, rather than on comparisons among individual lineages or on the PCA alone.

**Figure 3.**
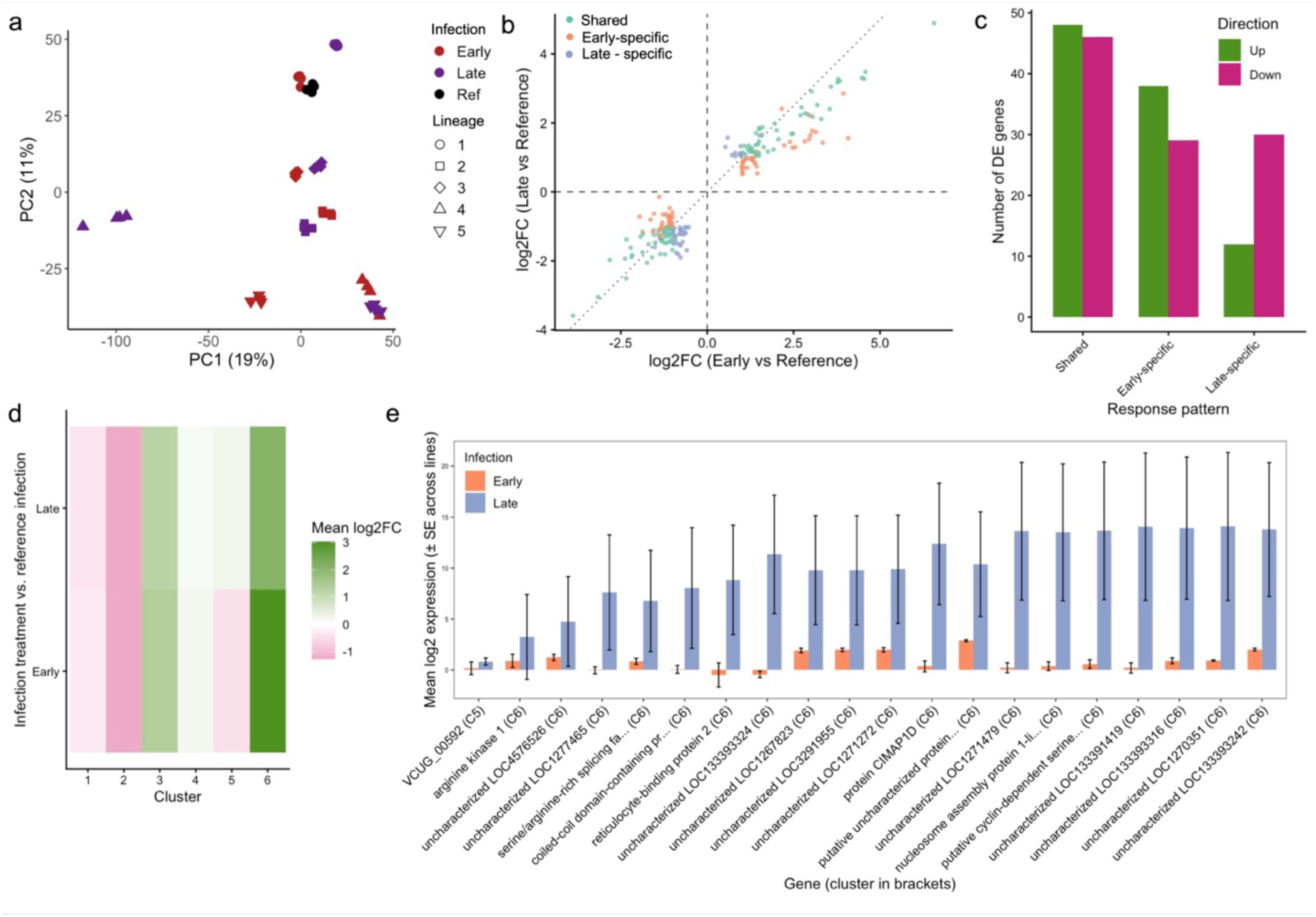
Parasite evolution reshapes the mosquito infection response. **(a)** PCA of infected treatments only, comparing mosquitoes infected with *reference* (**black**), *early* (**purple**) or *late* (**red**) parasite lineages. Each point represents one pool of ten females. Different symbols indicate the five independently evolved parasite lineages within the *early* and *late* treatments. PCA was performed on the 75% most variable genes across infected samples. *Early* and *late* infections both diverged from the *reference* response, but did so in different regions of PC space. **(b)** Comparison of log₂ fold-changes for *early vs. reference* (x-axis) and *late vs. reference* (y-axis). Each point represents one gene. Genes that were differentially expressed (FDR < 0.05 and |log₂FC| > 1) in both contrasts with the same sign are shown as shared (**green**), whereas genes that were differentially expressed only in *early* (**orange**) or only in *late* (**blue**) are shown as treatment-specific. Dashed lines indicate zero log₂FC, and the dotted diagonal indicates equal responses in *early* and *late*. **(c)** Numbers of up-regulated (**green**) and down-regulated (**pink**) genes in each response category from panel (b): shared, *early*-specific and *late*-specific.**(d)** k-means clustering of genes that were DE in at least one comparison involving *reference* infections (*reference vs. uninfected*, *early vs. reference*, late *vs*. *reference*). Clustering was performed on triplets of log₂FC values (missing values set to zero), with k = 6 clusters. The heatmap shows mean log₂FC for *early vs. reference* and *late vs. reference* within each cluster, highlighting groups of genes that respond more strongly in *early* or *late* infections. **(e)** Mean log₂ expression (± SE across lines) of a subset of genes belonging to clusters 5 and 6, which showed particularly strong divergence between *early* and *late* infections. Expression was averaged across replicate pools within each parasite lineage and then across the five *early* or five *late* lineages.

Pairwise differential expression analyses against the *reference* infection showed that both *early* and *late* infections altered the mosquito transcriptome, with a subset of genes similarly regulated in both conditions and additional sets uniquely affected in either *early* or *late* infections **(Fig. 3b, c; Supplementary Data 2)**. Genes were classified based on whether they were differentially expressed (FDR < 0.05; |log₂FC| > 1) in *early vs. reference*, in *late vs. reference*, in both contrasts in the same direction, or in neither. A substantial number of genes were regulated in the same direction in both *early* and *late* infections relative to the *reference*, whereas distinct sets of genes were specific to *early* or to *late* infections **(Fig. 3c)**. These comparisons are based on expression differences estimated across technical replicate pools for each lineage. Because measurements were taken at a single time point, these differences may also reflect variation in parasite load, infection dynamics, or response timing.

To summarise patterns across comparisons, genes that were differentially expressed in at least one of the three infection contrasts (*reference vs. uninfected*, *early vs*. *reference*, *late vs*. *reference*) were clustered by their log₂ fold-change profiles. Six k-means clusters captured distinct trajectories of how gene expression changed across the infection sequence **(Fig. 3d; Supplementary Data 3)**. Some clusters contained genes induced by the *reference* infection and showing higher expression in *early* infections but lower expression in *late* infections, whereas others showed the opposite pattern, with higher expression in *late* infections. These clusters, therefore, group genes with similar response profiles across parasite strategies but, on their own, do not indicate the functional processes involved and are used here to summarise consistent differences in expression patterns prior to functional analyses. Differences between *early* and *late* infections were consistent across multiple genes within each cluster **(Fig. 3e)**.

### Co-expression modules link infection backgrounds to eco-evolutionary traits

WGCNA was used to group genes into co-expression modules based on their expression profiles across all samples and to relate these modules to infection backgrounds and independently measured life-history traits. Seventeen modules (M0–M16) were detected. Correlations between module eigengenes and infection treatments showed that several modules tracked specific infection backgrounds **(Fig. 4a; Supplementary Data 4 and 5)**. Modules M14, M15, M6 and M9 were positively associated with *early* infections. In contrast, modules M0 and M16 were positively associated with *late* infections. These modules, therefore, group genes with expression patterns that differ across parasite infection strategies within the shared mosquito background.

**Figure 4.**
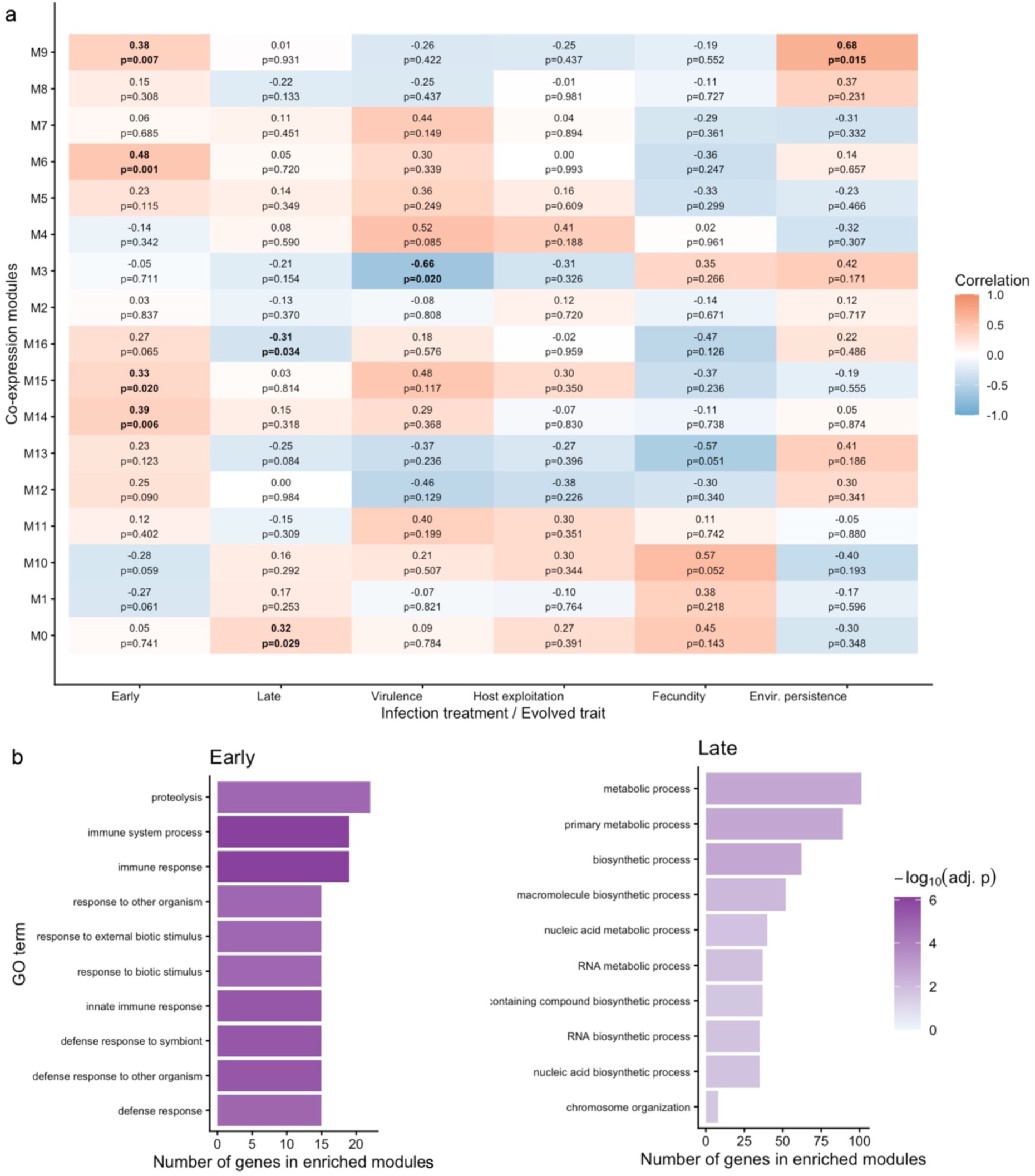
Co-expression modules connect evolved parasites with immune and metabolic gene programmes. **(a)** Correlations between module eigengenes and infection backgrounds or evolved life-history traits. Rows correspond to WGCNA modules (M0–M16). Columns correspond to infection treatments by evolved parasites (*early* or *late*) or line-level traits: virulence (maximum mortality hazard), host exploitation (spore density at day 10), fecundity at day 10, and environmental persistence (infection probability after 90 days outside the host). Each tile shows the Pearson correlation coefficient (colour scale) and the associated *p*-value; significant associations (p < 0.05) are indicated in bold. Modules M14, M15, M6 and M9 showed positive associations with *early* infections, whereas M0 and M16 were positively associated with *late* infections. Module M3 was negatively correlated with virulence, and module M9 was positively correlated with environmental persistence. **(b)** Gene Ontology (GO) enrichment of modules linked to *early* and *late* infections. Modules most strongly associated with *early* infections (M14, M15, M6 and M9; left panel) or *late* infections (M0 and M16; right panel) were grouped, and GO Biological Process enrichment was tested relative to the expressed gene universe. Bars show the number of genes in the enriched module set that mapped to each GO term, ordered by gene count. Bar colour intensity represents –log₁₀ of the adjusted *p-*value. *Early*-biased modules were enriched for immune and defence processes, whereas *late*-biased modules were enriched for RNA metabolic and core biosynthetic processes.

The same modules were then related to virulence (host mortality), host exploitation (parasite growth), fecundity (egg production) and parasite environmental persistence (spore infectivity after environmental exposure), previously measured in the same evolved lines [1,48]. Module M3 was negatively correlated with virulence, such that lineages with higher M3 expression showed lower mortality hazards. In contrast, module M9 was positively correlated with environmental persistence, indicating that higher expression of this module was associated with parasite lines whose spores remained infective for longer outside the host. These correlations do not imply a direct causal effect of host gene expression on parasite traits. Instead, they indicate that variation in host transcriptional state is associated with parasite traits measured across lineages. In the case of environmental persistence, this association may reflect differences in the host environment during spore production or release, rather than a direct effect on spore survival after release. Modules associated with *early* or *late* infections showed weaker or more trait-specific correlations, indicating that only a subset of infection-associated modules was strongly linked to measured life-history traits.

Functional enrichment of infection-linked modules indicated differences in the biological processes represented across modules. *Early*-associated modules (M14, M15, M6, and M9) were enriched for immune- and defence-related Gene Ontology (GO) terms, including responses to pathogens, proteolysis, and signalling processes **(Fig. 4b; Supplementary Data 6)**. In contrast, *late*-associated modules (M0 and M16) were enriched for RNA metabolic processes and core biosynthetic functions. These results show that modules associated with *early* and *late* infections differ in their enrichment for immune- and metabolism-related functions. Because enrichment analyses are based on overrepresented gene categories, these differences do not imply a strict functional partitioning of the transcriptome.

### Highly connected genes in modules associated with virulence and environmental persistence

To characterize the composition of modules associated with virulence and environmental persistence, intramodular connectivity (kME) was used to rank genes based on their correlation with module eigengenes. Modules that were correlated with virulence and environmental persistence were examined in more detail. In module M3, which was negatively correlated with virulence, genes were ranked by kME, and the highest-ranked genes are shown to illustrate module composition (n = 25). A similar set of highest-ranked genes is shown for module M9, which was positively correlated with environmental persistence. These genes represent highly connected members of modules associated with the measured traits and are shown to illustrate the expression patterns within these modules, but do not imply direct functional links to those traits.

Heatmaps of mean expression of these genes across the four infection treatments show distinct patterns for M3 and M9 **(Fig. 5; Supplementary Data 7)**. Genes in M3 tend to show expression profiles consistent with the negative association between this module and virulence, whereas genes in M9 show patterns consistent with the positive association with environmental persistence. These patterns reflect coordinated shifts in host transcriptional states across infection regimes rather than gene-level mechanisms.

**Figure 5.**
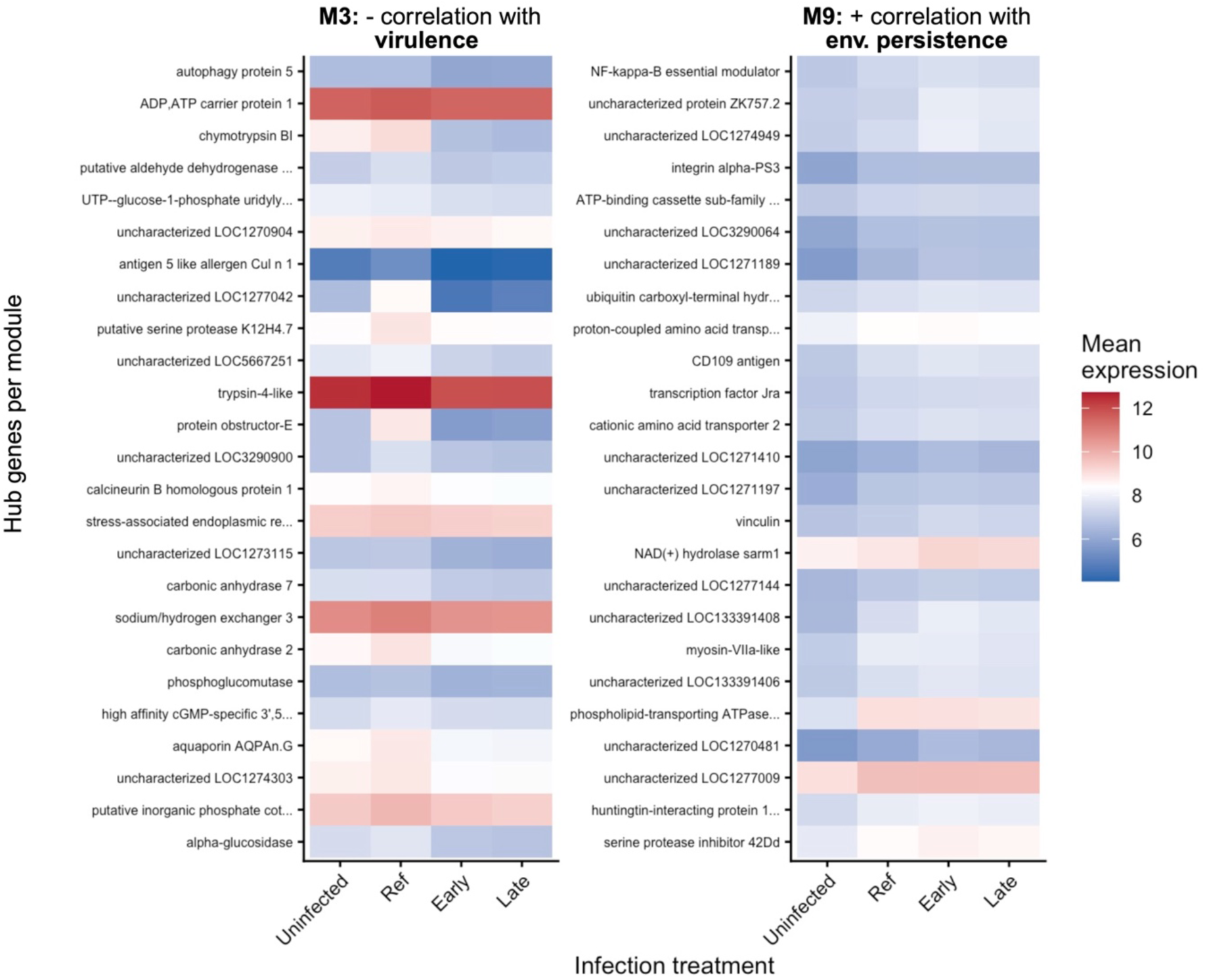
Highly connected genes in virulence- and environmental persistence-associated modules show distinct expression profiles across infection regimes. Heatmaps of mean expression for the top 25 highly connected genes (highest intramodular connectivity, kME) in module M3 (left) and module M9 (right). Module M3 showed a negative correlation with virulence, whereas module M9 showed a positive correlation with environmental persistence (Fig. 4a). Rows correspond to individual highly connected genes, ordered within each module by decreasing |kME|. Columns correspond to infection treatments: *uninfected*, *reference*, *early* and *late*. For each gene × treatment combination, normalized expression values were transformed to log₂(count + 1) and averaged across all replicate samples belonging to that treatment. Colour scale indicates mean expression, with red tones denoting higher expression and blue tones denoting lower expression. Together, these patterns illustrate how highly connected genes within each module vary across infection regimes and summarise the expression structure of modules associated with virulence and environmental persistence.

## Discussion

Parasites employ a wide range of strategies and life cycles, many of which are shaped by their ecology, including trade-offs between within-host performance and survival from one host to another. Evolutionary theory predicts that these differences in parasite strategy should be expressed through the host, as variation in exploitation, per-parasite pathogenicity, resistance, and tolerance jointly determines virulence, here defined as infection-induced host mortality [2,6,18]. In this system, lineages of the microsporidian *V. culicis* have been selected for earlier or later transmission, generating parasite populations that differ in growth, virulence and environmental persistence while sharing the same *An. gambiae* host background. Previous work on these lineages has shown that *late*-selected parasites grow faster, reach higher spore loads, and are more virulent, whereas *early*-selected lines show lower exploitation and virulence but produce spores that remain infective for longer outside the host [1,47,48]. The present study adds a transcriptomic layer at a single, common time point (day 10 of adulthood) to examine how these evolved parasite life histories are associated with host gene expression and how host co-expression modules covary with virulence, host exploitation, fecundity, and environmental persistence **(Fig. 1)**. The analyses presented here identify host transcriptional states that covary with parasite life-history traits, rather than causal mechanisms linking specific genes to infection outcomes. Because gene expression was measured at a single time point, the results represent a stage-specific snapshot of host–parasite interactions.

Within this snapshot, variation among parasite lineages provides an additional axis of biological structure. The clustering of samples by parasite lineage indicates that lineage-level (or selection replicate-level) differences contribute strongly to transcriptomic variation, consistent with independently evolving populations following distinct trajectories under the same selection regime.

### A conserved core response to infection provides the baseline

The comparison between *uninfected* and *reference*-infected mosquitoes revealed a strong and consistent infection signature **(Fig. 2)**. Samples separated clearly in multivariate space, and hundreds of genes were differentially expressed. Previous work showed that reference infections already impose detectable costs on survival and fecundity by day 10 [1]. The core transcriptional response observed here at the same time point likely reflects parasite recognition, canonical immune effectors (including melanization, antimicrobial peptides, and cellular responses), and systemic adjustments in metabolism and stress management.

The spread among replicate pools observed in the PCA indicates that a substantial proportion of the variance in the dataset is not captured by infection status alone. This variability likely reflects multiple sources of biological heterogeneity. First, the mosquito population is not isogenic, and differences in host genotype or physiological state can influence infection outcomes and transcriptional responses [49,50]. Second, even under standardized exposure, infections can vary in intensity and progression, leading to differences in parasite load and host condition at the time of sampling [51]. Third, independently evolving parasite lineages may follow only partially parallel trajectories under the same selection regime, as expected in experimental evolution, where standing genetic variation, contingency, and drift can generate divergence among replicate lines [52–54]. Together, these sources of variation are consistent with the observed spread among samples and highlight that inference in this system should rely on consistent differences across treatments rather than on any single lineage or pool.

When this response is compared with an earlier study using *V. culicis* in *Aedes aegypti*, more genes and larger expression changes were reported in *Aedes* than in *Anopheles* [55]. This difference may reflect stronger or more extensive transcriptional responses in *Aedes*, potentially including immune-related processes [37]. In other words, *V. culicis* elicits a robust but host-specific programme, and the *Anopheles* response appears more constrained. This *reference* response in *An. gambiae* provides a valuable baseline against which divergence in host responses due to parasite life-history evolution can be interpreted.

### Divergent host responses under early and late infection

When only infected treatments were considered, *early*, *late* and *reference* infections occupied overlapping but distinct regions of transcriptomic space **(Fig. 3a)**. Variation among samples was structured both by parasite lineage and by infection treatment, with lineage-level differences contributing strongly to the overall spread. All infections shared the core response, consistent with the fact that by day 10, all lines had completed their life cycle and were producing spores [1]. However, pairwise comparisons against the reference revealed a mixture of shared and regime-specific changes. Many genes were differentially expressed in both *early* and *late* infections in the same direction, whereas others changed only in *early* or only in *late* lines **(Fig. 3b, c)**. These treatment-level differences are observed consistently across independent lineages, indicating that they are not driven by any single lineage but reflect reproducible differences between parasite strategies.

Clustering log₂ fold-changes across *reference vs. uninfected*, *early vs. reference* and *late vs. reference* summarised these patterns into a small number of shared and divergent expression patterns across infection conditions **(Fig. 3d)**. Some clusters represented a generic infection response that was further amplified by both evolved regimes whereas others contained genes whose expression differed between *early* and *late* infections at the same time point **(Fig. 3e)**. Because all measurements were taken at a single time point (day 10), these differences reflect variation observed at the sporulating stage of infection but may also be influenced by differences in infection dynamics, including parasite load or timing. Distinguishing between these possibilities would require temporal sampling across infection stages. Such experiments are considerably more resource-intensive in this system, and the present study provides an initial framework for identifying whether broader temporal monitoring is likely to be informative.

These patterns are consistent with the broader life-history phenotyping. *Late* parasites grow faster, reach higher chronic loads, cause higher adult mortality and induce earlier reproduction and shifts in resource allocation than early parasites [1,47,48]. The separation of *early* and *late* infections across multiple gene clusters, therefore, indicates consistent stage-specific differences in host gene expression associated with parasite strategy.

### Co-expression modules link infection backgrounds to host response space

The co-expression network analysis provides a more integrated view of these differences **(Fig. 4)**. WGCNA grouped genes into modules whose eigengenes varied across *uninfected*, *reference*, *early* and *late* infection treatments. Some modules behaved as general infection modules, with elevated eigengenes in all infected treatments relative to *uninfected* controls. Others were more clearly associated with *early* or *late* infections, indicating that experimental evolution has altered how specific gene networks are regulated.

Modules most strongly associated with *early* infections (M14, M15, M6, and M9) were enriched for immune, defence, and stress-related GO terms, as well as transcriptional and regulatory processes. In contrast, modules associated with late infections (M0 and M16) were enriched for RNA metabolism, biosynthesis and broader metabolic functions. Although GO categories are coarse, this separation suggests that *early* and *late* parasite regimes push the host into different regions of response space. *Early* parasites, which are less virulent and reach lower chronic loads, are likely still constrained by classical immune and stress responses reflected in these modules. For *late* parasites, which are more virulent and reach higher loads, immune-annotated modules are less prominent; instead, transcriptional and metabolic rewiring dominate.

Several non-exclusive mechanisms could underlie this pattern. *Late* parasites may reduce early immune recognition or dampen effector deployment [56–58], allowing faster growth despite a comparatively “quieter” immune readout. Hosts may rely more on metabolic reorganization and damage control to maintain key functions at higher parasite burdens, a pattern closer to disease tolerance than to resistance. In addition, independent life-history work showed that *late* infections induce earlier reproduction and a shift away from somatic maintenance, consistent with terminal investment by the host under high mortality risk [33,34]. Taken together, the module structure suggests that *early* infections are characterized by a stronger immune and stress response, whereas *late* infections are characterized by greater changes in biosynthesis, energy use, and metal handling.

These module-level patterns align closely with previous biochemical and metallomics phenotyping of the same lines. *Late* infections increased protein and carbohydrate reserves and altered metal homeostasis, especially for iron, zinc and manganese, relative to *reference* infections [47]. Modules that are upregulated in *late* infections and enriched for metabolic and metal-related functions, therefore, provide a plausible molecular counterpart to these organismal changes.

### A host expression module associated with lower virulence and consistent with disease tolerance

A central aim of this study was to relate host transcriptional variation to the eco-evolutionary outcomes previously quantified in these parasite lineages [1,47,48]. Among the co-expression modules, M3 showed the clearest association with virulence, with higher module activity consistently observed in infection backgrounds exhibiting lower mortality hazards. This relationship was robust across independently evolved parasite lineages, indicating that variation in host gene expression covaries with evolved differences in infection outcomes. This pattern suggests that parasite evolution consistently aligns with alternative host expression states, rather than with isolated gene-level changes.

The genes contributing most strongly to M3 were enriched for functions related to tissue maintenance, metabolic regulation, and cellular homeostasis, including components involved in autophagy, membrane integrity, ion transport, and epithelial structure. Notably, several highly connected genes in this module have previously been implicated in stress responses and tissue integrity in insects, including genes associated with autophagic processes, epithelial barrier maintenance, and metabolic regulation (e.g., Atg family genes, ion transporters, and cuticle-associated proteins) [10,59]. These functions are more readily interpreted as contributing to the maintenance of host function during infection, defining a host state in which damage is limited without necessarily reducing parasite burden, consistent with processes related to disease tolerance [8,15,16,60–63].

Crucially, this interpretation is supported by alignment between module activity and independently measured virulence across replicate lineages, rather than by gene function alone. However, these associations remain correlational and may also reflect variation in parasite load or infection dynamics.

Interpreted in this context, M3 defines a candidate host response linking parasite strategy to variation in infection outcomes. Such responses provide a tractable starting point for future functional work aimed at disentangling resistance, tolerance, and other mechanisms underlying virulence in this system.

### A host expression module associated with environmental persistence

In contrast to M3, module M9 showed the strongest association with parasite environmental persistence, with higher module activity observed in infection backgrounds whose spores remained infective for longer outside the host. This association was consistent across independently evolved parasite lineages, indicating that host transcriptional variation covaries with variation in parasite life-history traits measured in the same system.

Genes with high connectivity within M9 were enriched for functions related to membrane dynamics, signalling, lipid handling, and cellular organization, including processes involved in vesicle trafficking, cell-surface structure, and metabolic regulation. Several of these genes are linked to pathways involved in secretion, membrane remodelling, and host–parasite interface dynamics in insects, including vesicle trafficking components, lipid metabolism genes, and membrane-associated signalling proteins. These functions are compatible with pathways that define the physiological and structural environment in which parasite transmission stages develop within the host.

The positive association between M9 activity and environmental persistence is therefore consistent with variation in host physiological states at the point of transmission, suggesting that host environments may influence the quality of transmission stages produced [3,64–66]. However, these associations remain correlative and may also reflect variation in parasite traits or infection dynamics.

Within this context, M9 defines a candidate host response associated with parasite persistence beyond the host. This identifies a testable hypothesis that the host state at the point of transmission contributes to variation in parasite persistence across environments, linking within-host processes to transmission outcomes.

Several limitations should be considered when interpreting these results. First, gene expression was measured at a single sporulating stage of infection, and observed differences may therefore reflect variation in parasite load, infection dynamics, or response timing rather than entirely distinct transcriptional programmes. Second, the analyses identify associations between host transcriptional states and parasite life-history traits, but do not establish direct causal relationships between specific genes and infection outcomes. Finally, independently evolving parasite lineages showed substantial variation, indicating that partially distinct evolutionary trajectories can emerge under similar selection regimes. Despite these limitations, consistent differences across early and late treatments support the conclusion that parasite life-history evolution is associated with reproducible changes in host transcriptional responses.

## Conclusions

Overall, this study links experimental evolution, life-history phenotypes and host transcriptomic architecture in a natural mosquito–microsporidian system. *Early* and *late* parasite lines differ consistently in life-history traits, with *late* lines showing higher parasite growth, higher virulence (infection-induced mortality), and reduced environmental persistence, whereas *early* lines show lower virulence and longer persistence of transmission stages. It shows that evolved parasite life history is reflected in consistent, stage-specific differences in host gene expression, identifies a co-expression module consistent with disease tolerance, and highlights a second module associated with environmental persistence. These results indicate that associations between virulence, tolerance, host exploitation, and environmental persistence can be captured at the level of host co-expression modules, providing a framework for linking parasite life-history evolution to host biology.

Taken together, these modules have broader implications for microbial ecology, disease ecology and epidemiology. The tolerance-associated M3 module is consistent with host processes that limit damage without necessarily reducing parasite load. In contrast, the environmental-persistence module M9 is associated with parasite lines whose spores remain infective for longer outside the host. These associations suggest that host physiological state at the time of spore production and release may influence the conditions under which transmission stages are produced, rather than directly determining their survival after release. In this sense, hosts may contribute to shaping the environments encountered by parasite transmission stages, although the mechanisms underlying these links remain to be established.

More broadly, the approach used here, *i.e.* evolving parasite life histories and analyzing the resulting host transcriptional responses, can be applied across vector and non-vector systems to dissect how resistance, tolerance and transmission are implemented in natural host-parasite systems. Importantly, this study shows that evolutionarily-grounded descriptive, system-level analyses are a necessary step before mechanistic or functional work (e.g., functional genetics), as they identify the axes of variation and candidate processes that structure infection outcomes. Extending this framework to additional host–parasite combinations, multiple time points, and more complex environments (e.g., with seasonality) will help clarify how host processes shape the environments parasites experience and influence infection dynamics and evolution.

## Materials and Methods

### Model system

The Kisumu strain of the mosquito *Anopheles gambiae (s.s.)* was used as host. As parasites, 11 independently maintained lineages of the microsporidian *Vavraia culicis floridensis* [36,37] were used: one unselected laboratory population (*reference*), five lineages selected for early transmission (*early*) and five selected for late transmission (*late*). *Early* lineages were obtained by repeatedly collecting spores from the earliest dying hosts in each generation, favouring parasites that completed their life cycle rapidly and killed hosts sooner. *Late* lineages were obtained by collecting spores from the latest dying hosts, favouring parasites that took longer to kill their hosts and therefore had a later transmission [1]. The performance of these lineages and their effects on mosquito life history, virulence and parasite exploitation have been characterized in detail [1,36,46,47,67]. All experiments were conducted under standard conditions (26 ± 1 °C, 70 ± 5% relative humidity, 12:12 h light-dark cycle).

### Infection assay and sample collection

Freshly hatched *An. gambiae* larvae were reared in Petri dishes at a density of 50 larvae in 80 ml of distilled water and fed daily with fish food according to a standard regime [1]. On day 2 of larval development, individuals were exposed to 10,000 *V. culicis* spores per larva from the respective parasite line (*early*, *late* or *reference*); *uninfected* controls were handled identically but received no spores. At pupation, individuals were transferred to tubes containing distilled water and allowed to emerge. Adult females were maintained with constant access to 6% sucrose solution.

Females were sampled on day 10 of adulthood, when previous work on these lines has shown that all infections produce infective spores and that virulence differences among parasite lineages are expressed [1]. On that day, all surviving females were collected and stored at −80 °C until RNA extraction. For RNA sequencing, each parasite lineage (*reference*, five *early* lines and five *late* lines; n = 11) and the *uninfected* control were represented by four independent biological replicates, each consisting of an independent pool of 10 females. Replication, therefore, occurred at two levels: each lineage (and the *uninfected* control) was sampled four times, and the *early* and *late* treatments each comprised five independently evolved parasite lineages.

### RNA extraction, library preparation and sequencing

Pools of 10 females were homogenized in QIAzol reagent with stainless steel beads using a TissueLyser LT (30 Hz, 3 min), and RNA was extracted using the RNeasy Universal Plus kit (Qiagen), including genomic DNA removal according to the manufacturer’s protocol. Individuals were randomly assigned to pools. Adults were homogenized in QIAzol reagent with stainless steel beads, and RNA was eluted in nuclease-free water. RNA concentrations were adjusted before shipment on dry ice to the Genomic Technologies Facility at the University of Lausanne (Switzerland). RNA quality was assessed on a Fragment Analyzer. Libraries were prepared from 250 ng total RNA per sample using the Illumina Stranded mRNA Prep kit with unique dual indices, quantified and checked for size distribution, and sequenced as single-end 150-bp reads on an Illumina NovaSeq 6000 platform, yielding approximately 20–38 million reads per sample after demultiplexing.

### Read processing and gene quantification

Adapters and low-quality bases were removed with Cutadapt, and short reads (< 40 nt) were discarded. Reads mapping to rRNA were removed with fastq_screen. Filtered reads were aligned against a concatenated reference combining the *An. gambiae* genome (assembly GCA_943734735.2) and the *V. culicis floridensis* genome (GCA_000192795.1) using STAR. Gene-level counts were obtained with htseq-count using the corresponding GTF annotation files. Alignment quality was assessed with RSeQC; libraries showed >93% alignment to the mosquito genome and broad coverage across expressed genes. Analyses focus on host (*An. gambiae*) gene expression; parasite reads were retained during alignment but were not analyzed further here **(Supplementary Table 1, Supplementary Fig. 1)**.

### Expression filtering and normalization

Bioinformatic processing and downstream analyses were conducted in R (version 4.3.1). Gene-level count tables were filtered to remove lowly expressed genes, retaining genes with counts per million (CPM) greater than 1 in at least 4 libraries. Library sizes were normalized using the trimmed mean of M-values (TMM) method. Log₂-counts-per-million (logCPM) values were calculated using a prior count (pseudocount) of 2, as implemented in edgeR, to stabilize variance for lowly expressed genes. These logCPM values were used for exploratory analyses and linear modelling. For multivariate and network analyses, variance-stabilized expression measures derived from the normalized counts were used where appropriate. The normalized expression matrix was saved as a genes × samples table and imported into R for the statistical analyses described below.

### Statistical analyses

Analyses and plotting were performed in R 4.3.1 using standard tidyverse packages [68], as well as clusterProfiler [69] and WGCNA [70]. Analyses were structured to (i) quantify a core infection signature by comparing *reference* infections with *uninfected* controls; (ii) quantify divergence in host expression between *early* and *late* infections relative to the *reference*; and (iii) identify co-expression modules and relate them to independent measurements of parasite life-history traits (virulence, host exploitation, fecundity and environmental persistence) obtained from the same evolved lines.

For principal component analysis (PCA), the upper 75% most variable genes were retained to reduce noise from low-variance genes while preserving the majority of biologically relevant variation; similar qualitative structures were observed across nearby thresholds (e.g., 50–90%), indicating that the choice does not affect the main conclusions. PCA was then performed on the transposed, centred and scaled expression matrix using prcomp [71].

Differential expression between treatments was assessed using gene-wise two-sample t-tests on normalized log₂-transformed expression values. Log₂-counts-per-million values approximate normality after transformation, allowing the use of linear models on gene-wise expression, and the large number of genes analyzed enables robust multiple testing correction. This approach provides a transparent framework for identifying large, consistent expression differences, and the main conclusions are supported by effect sizes and consistency across independent parasite lineages. P-values were adjusted using the Benjamini–Hochberg correction.

Unless stated otherwise, genes with a false discovery rate (FDR) < 0.05 and an absolute log₂ fold-change > 2 were considered differentially expressed. This threshold was used to identify robust expression changes for primary comparisons, whereas a more permissive threshold (FDR < 0.05; |log₂ fold-change| > 1) was used in overlap and clustering analyses to capture broader expression patterns and avoid excluding genes with moderate but consistent changes across conditions.

To analyze patterns across infection treatments, log₂ fold-changes from the three contrasts (*reference vs. uninfected*, *early vs*. *reference*, *late vs. reference*) were combined. Genes that were differentially expressed in at least one contrast were retained, and k-means clustering (k = 6) was applied to their log₂ fold-change profiles to group genes with similar expression patterns across conditions. The choice of k = 6 provided a balance between capturing major expression patterns and avoiding over-fragmentation; similar qualitative clustering structures were observed across nearby values of k, indicating that the results are not sensitive to this parameter choice.

Weighted gene co-expression network analysis (WGCNA) was used to identify modules of co-regulated genes and relate them to infection treatment and evolved life-history traits. A signed network was constructed from variance-filtered and normalized expression values. Modules were defined using dynamic tree cutting with a minimum module size of 30 genes and a merge cut height of 0.25. The soft-thresholding power (β) was selected based on the scale-free topology criterion, following standard WGCNA guidelines, and a value of 6 was chosen as the lowest power at which the scale-free topology fit index reached a plateau while maintaining sufficient mean connectivity [70].

Module eigengenes were correlated with binary indicators for each infection treatment and with line-level trait means for virulence, host exploitation, fecundity and environmental persistence. Gene Ontology (GO) enrichment for module subsets associated with early and late infections was tested using enrichGO with the “org.Ag.eg.db” annotation [72], defining the expressed gene set as the background. Within the WGCNA network, intramodular connectivity (kME) was used to identify highly connected genes in modules linked to virulence (M3) and environmental persistence (M9). Gene annotations were obtained from VectorBase and OrthoMCL-DB [73,74].

## Author contributions

LMS designed and conducted the experiments, analyzed the data and wrote the manuscript.

## Acknowledgements

I thank Amjad Khalaf, Ian Will, Paula Escuer Pifarré, Thomas J. Travers-Cook and Tiago G. Zeferino for their advice and experimental support. I also thank the Lausanne Genomic Technologies Facility (University of Lausanne, Switzerland) team, including Johann Weber, Julien Marquis, Sandra Calderon Copete, and Viviane Praz, for their assistance with library preparation, sequencing, and preliminary data analyses. I would especially like to thank Jacob C. Koella and Kayla C. King for their constant feedback, support, and critical comments on the first draft of the manuscript. The project was supported by the Basler Stiftung für biologische Forschung and LMS, funded by a SNF grant 310030_192786.

## Competing interests

The author declares that they have no competing interests.

## Data availability

Raw and processed RNA-sequencing data have been deposited in GEO under accession ID GSE336387. Additional datasets and R script code are available in Borealis Dataverse at https://doi.org/10.5683/SP4/O8LPID.

## Notes

### Competing Interest Statement

The authors have declared no competing interest.

### Summary of Updates

Last changes to the manuscript before publication.

