## Supplementary information for "Host transcriptional signatures associated with disease tolerance and environmental persistence in a mosquito–microsporidian system"

**Supplementary files index:**

**Supplementary information:** Supplementary tables and figures. (*current one*)

*Accompanying the manuscript:*

**Supplementary Data 1:** Differential expression (DE) between the *reference* and the *uninfected*.

**Supplementary Data 2:** DE between *early* and *late*, compared to the *reference.*

**Supplementary Data 3:** k-means reaction norms clusters.

**Supplementary Data 4:** WGCNA gene-module membership.

**Supplementary Data 5:** WGCNA module-trait correlations.

**Supplementary Data 6:** Gene ontology (GO) enrichment for *early*- and *late*-biased modules.

**Supplementary Data 7:** Hub genes in modules 3 and 9 and their expression.

*Deposited in GEO:*

**FASTQ files:** Raw RNA-sequencing data.

*Deposited in Borealis dataverse:*

**File S10.** Virulence data (and the respective readme file).

**File S11.** Fecundity data (and the respective readme file).

**File S12.** Environmental persistence data (and the respective readme file).

**File S13.** R script for statistical analyses and plots.

This document contains the following:

**Table S1.** Number of genes detected per sample.

**Table S2**. Principal component analysis (PCA) of the host response to the *reference* infection compared to *uninfected* hosts.

**Table S2**. Principal component analysis (PCA) of the host response to the *early* and *late* infection compared to *reference* hosts.

**Figure S1.** Host RNA-sequencing and parasite gene counts.

**Figure S2.** Quality control clustering and PCAs.

**Figure S3.** Heatmap of the log-fold change for all transcripts across all biological replicates.

**Figure S4.** Volcano plots of the host response to *early*- and *late*-infections compared to the *reference* one.

**Tables:**

**Table S1. Number of genes detected per sample.** The number of known *Anopheles gambiae* genes per parasite lineage and biological replicate (four independent pools of ten females each) is listed below.

| Treatment | Lineage | Biological replicate | Sample name | Number of known genes detected |
|---|---|---|---|---|
| **Uninfected** | - | 1 | U11 | 9303 |
| | - | 2 | U12 | 9304 |
| | - | 3 | U13 | 9146 |
| | - | 4 | U14 | 9328 |
| Infected with **reference parasite (unselected)** | - | 1 | R11 | 8973 |
| | - | 2 | R12 | 8547 |
| | - | 3 | R13 | 8600 |
| | - | 4 | R14 | 8801 |
| **Infected** with parasites selected for **early-transmission** | 1 | 1 | E11 | 9433 |
| | 1 | 2 | E12 | 9043 |
| | 1 | 3 | E13 | 8859 |
| | 1 | 4 | E14 | 8847 |
| | 2 | 1 | E21 | 9113 |
| | 2 | 2 | E22 | 8557 |
| | 2 | 3 | E23 | 9017 |
| | 2 | 4 | E24 | 9282 |
| | 3 | 1 | E31 | 9122 |
| | 3 | 2 | E32 | 8984 |
| | 3 | 3 | E33 | 9020 |
| | 3 | 4 | E34 | 8901 |
| | 4 | 1 | E41 | 8799 |
| | 4 | 2 | E42 | 8982 |
| | 4 | 3 | E43 | 7962 |
| | 4 | 4 | E44 | 9070 |
| | 5 | 1 | E51 | 8807 |
| | 5 | 2 | E52 | 8951 |
| | 5 | 3 | E53 | 8851 |
| | 5 | 4 | E54 | 8784 |
| **Infected** with parasites selected for **late-transmission** | 1 | 1 | L11 | 9171 |
| | 1 | 2 | L12 | 9094 |
| | 1 | 3 | L13 | 8958 |
| | 1 | 4 | L14 | 9107 |
| | 2 | 1 | L21 | 9083 |
| | 2 | 2 | L22 | 9356 |
| | 2 | 3 | L23 | 9123 |
| | 2 | 4 | L24 | 8929 |
| | 3 | 1 | L31 | 8922 |
| | 3 | 2 | L32 | 9038 |
| | 3 | 3 | L33 | 9193 |
| | 3 | 4 | L34 | 9161 |
| | 4 | 1 | L41 | 8716 |
| | 4 | 2 | L42 | 8855 |
| | 4 | 3 | L43 | 8462 |
| | 4 | 4 | L44 | 7629 |
| | 5 | 1 | L51 | 9386 |
| | 5 | 2 | L52 | 9638 |
| | 5 | 3 | L53 | 9538 |
| | 5 | 4 | L54 | 9136 |

**Table S2. Principal component analysis (PCA) of the host response to the *reference* infection compared to *uninfected* hosts.** PCA variance for the analysis shown in Figure 2a (underlined values). Shown are the eigenvalues, proportion of variance, and cumulative variance explained by the first principal components describing variation in gene expression between Anopheles gambiae infected with *reference Vavraia culicis* or not (*uninfected*). These axes were used to summarise the core transcriptional response to infection that underlies subsequent comparisons with parasites evolved for *early*- or *late*-transmission.

| | Eigenvalue | Proportion of variance | Cumulative variance |
|---|---|---|---|
| **PC1** | <u>$2.50 \times 10^3$</u> | <u>0.336</u> | <u>0.336</u> |
| **PC2** | <u>$1.20 \times 10^3$</u> | <u>0.161</u> | <u>0.498</u> |
| **PC3** | $1.03 \times 10^3$ | 0.139 | 0.637 |
| **PC4** | $8.25 \times 10^2$ | 0.111 | 0.747 |
| **PC5** | $7.37 \times 10^2$ | 0.090 | 0.846 |

**Table S3. PCA of the host response to *early* and *late* infection compared to the *reference.*** PCA variance for the analysis shown in Figure 3a (underlined values). Shown are the eigenvalues, proportion of variance, and cumulative variance explained by the first principal components describing variation in gene expression between *Anopheles gambiae* infected with *early* or *late Vavraia culicis* or with the *reference* lineage.

| | **Eigenvalue** | **Proportion of variance** | **Cumulative variance** |
|---|---|---|---|
| **PC1** | <u>1387.51</u> | <u>0.186</u> | <u>0.186</u> |
| **PC2** | <u>835.98</u> | <u>0.112</u> | <u>0.299</u> |
| **PC3** | 387.21 | 0.052 | 0.351 |
| **PC4** | 371.36 | 0.050 | 0.401 |
| **PC5** | 343.87 | 0.046 | 0.447 |

**Figures:**

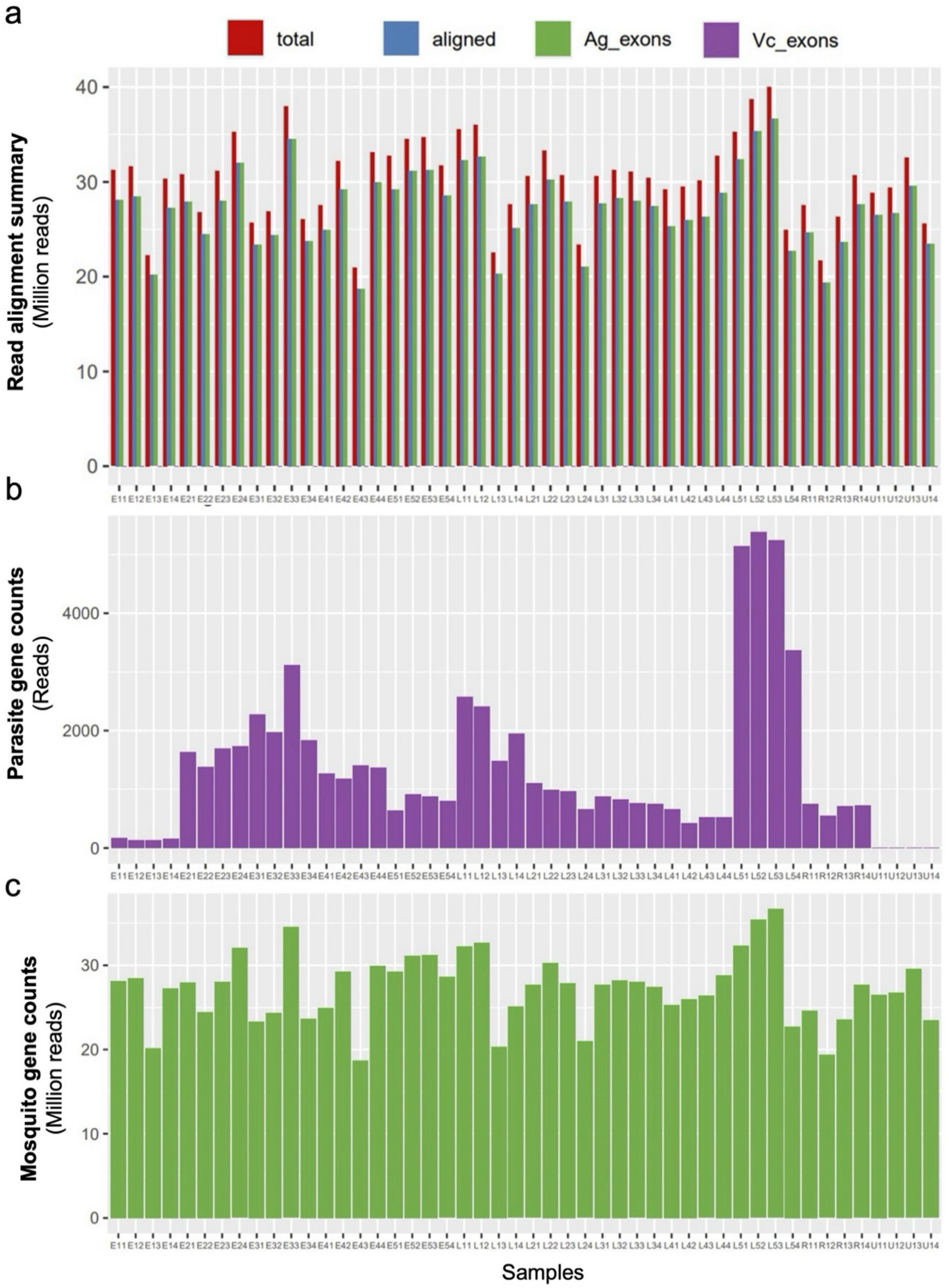


**Figure S1. Host RNA-sequencing and parasite gene counts. (a)** Barplot showing the number of total reads, the number of aligned reads, and how many of which are *Anopheles gambiae* or *Vavraia culicis* exons. **(b)** Number of *V. culicis* reads found in each of the samples. **(c)** Number of *A. gambiae* reads found in each sample.

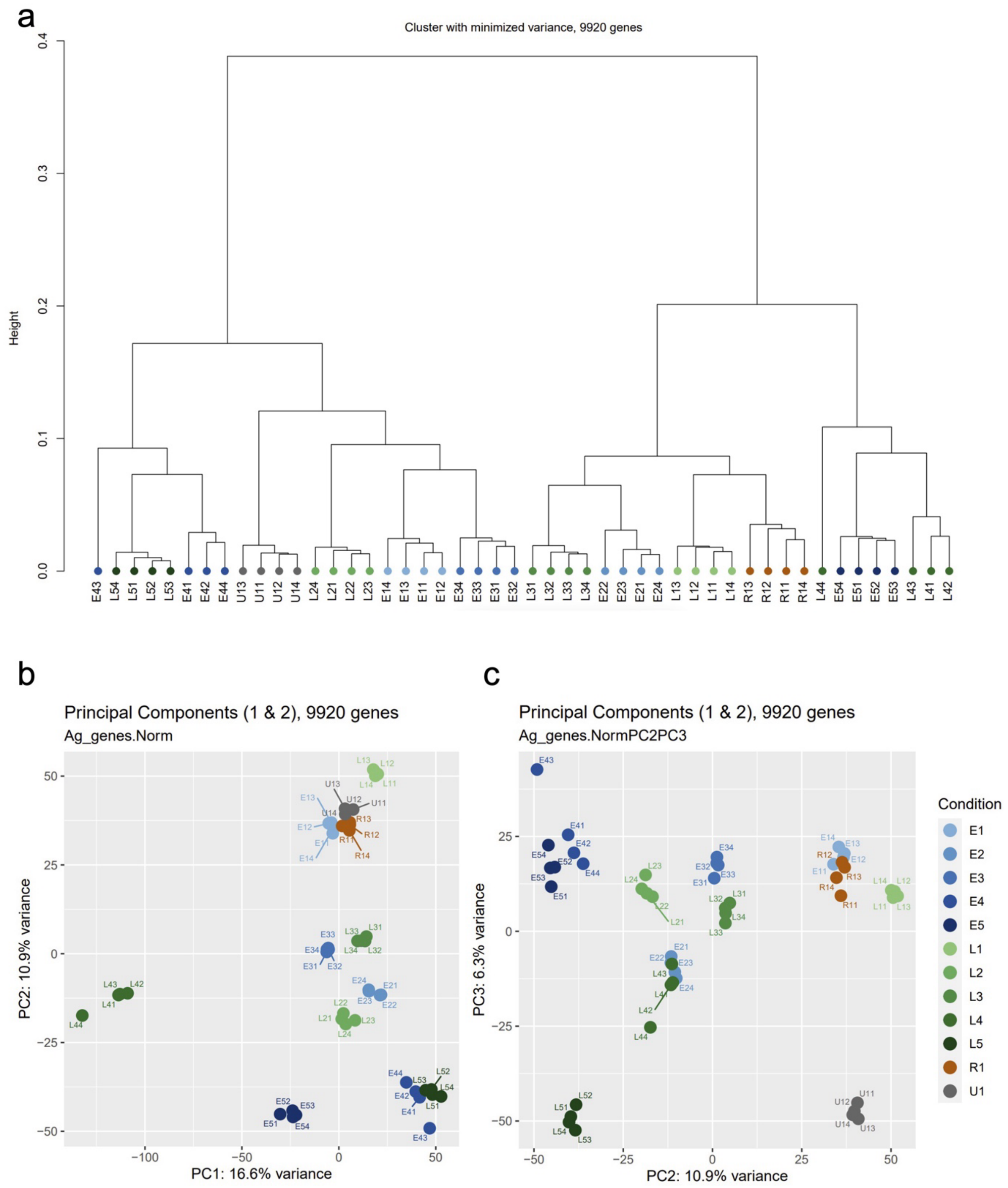


**Figure S2. Quality control clustering and PCAs.** Statistical quality controls were performed through density gene expression distribution, clustering, and sample PCA. **(a)** Clustering and **(b)** PCA show that replicates group together as expected, with two exceptions for samples E43 and L44. **(c)** Since the first two principal components in the PCA explain only 30% of the variability, a PCA including PC2 and PC3 (an additional 6% of the variability) was considered. The latter does not indicate a clustering of *uninfected* and unselected *reference* infections.

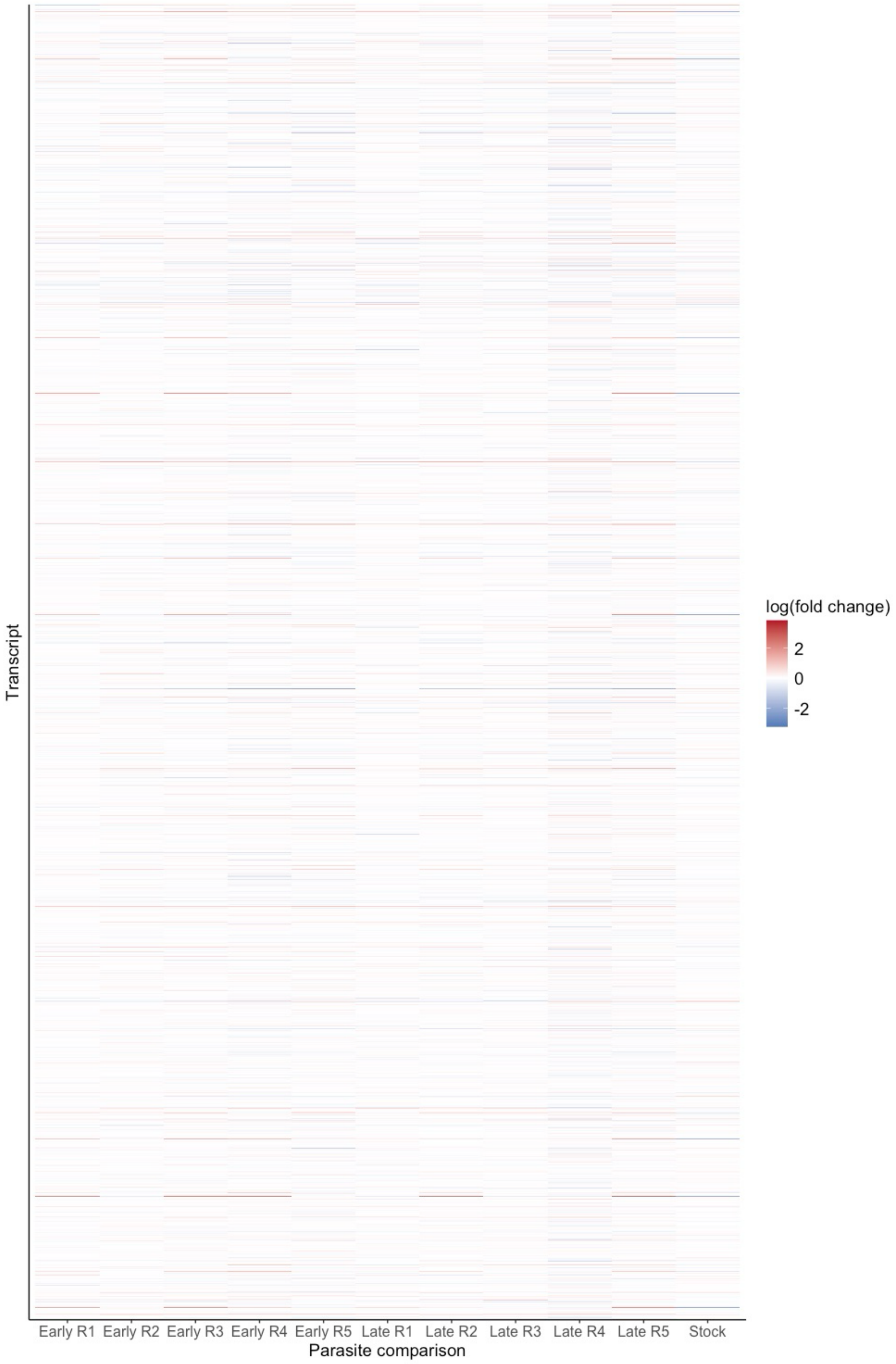


**Figure S3. Heatmap of the log-fold change for all transcripts across all biological replicates.** Heatmap showing the log-transformed fold-change in the expression of all detected transcripts (*i.e.,* 9920) by parasite treatment comparison. All host samples were compared with the host samples in response to the unselected *reference/stock* parasite, except for themselves, which were compared with the *uninfected* host transcriptome.

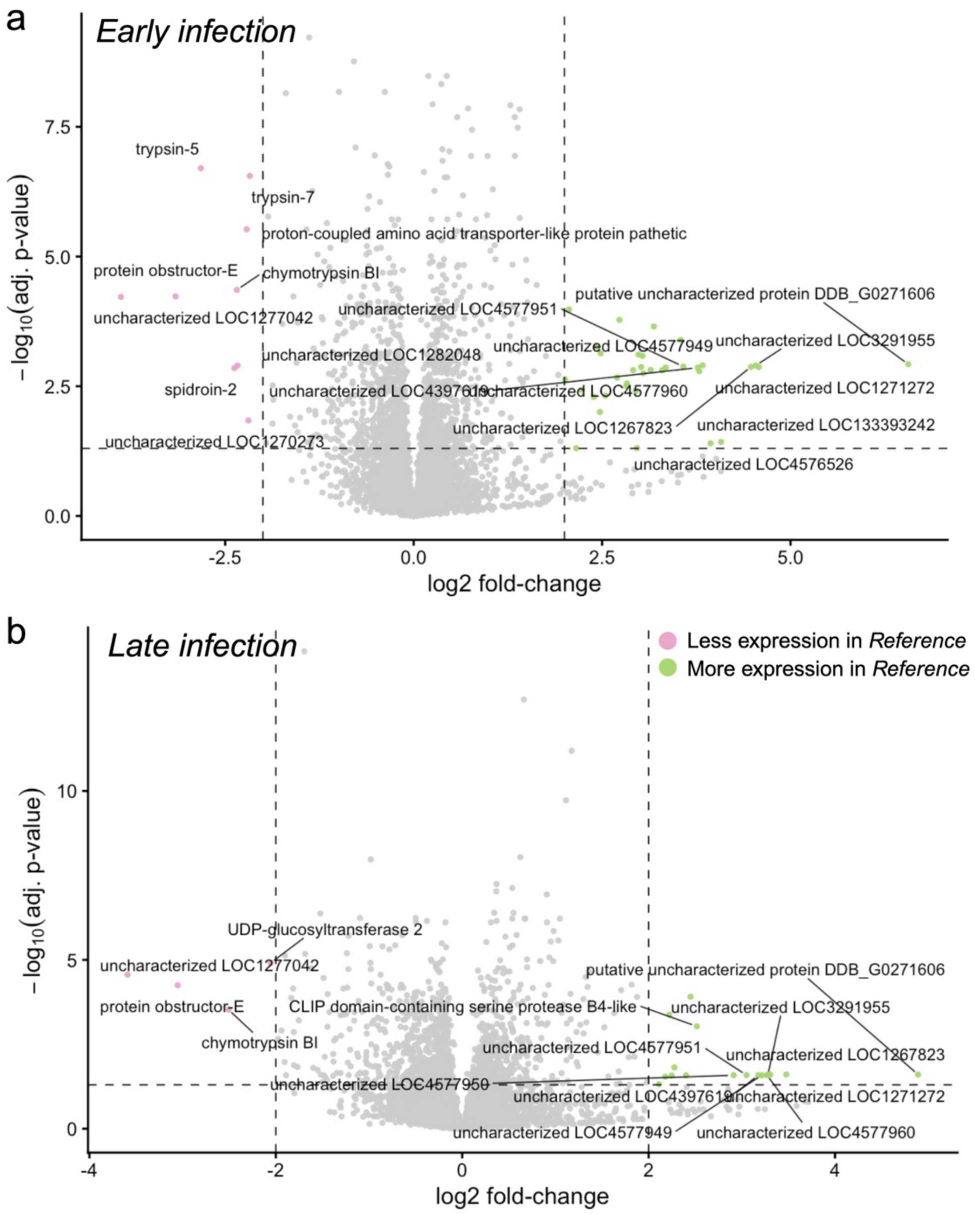


**Figure S4. Differential expression in *early*- and *late*-infections relative to the *reference* infection.** **(a)** Volcano plot of $\log_2$ fold-change *vs.* $-\log_{10}$(adjusted *p*-value) for *early vs. reference* infections. **(b)** Volcano plot for *late vs. reference* infections. Dashed vertical lines indicate the $|\log_2$ fold-change$|$ threshold ($\geq 2$), and the horizontal dashed line marks the FDR cutoff (adj. $p < 0.05$). Points in pink and green denote genes with significantly lower or higher expression, respectively, in the *reference* infection compared with the evolved infection; grey points are non–differentially expressed genes. Selected labelled genes highlight those with the largest and most significant changes.
